# Whatman FTA^®^ cards versus plasma specimens for the quantitation of HIV-1 RNA using two Real-Time PCR assays

**DOI:** 10.1101/748061

**Authors:** Abdourahamane Yacouba, Malika Congo, Gérard Komonsira Dioma, Hermann Somlaré, David Coulidiaty, Kalifa Ouattara, Lassana Sangare

## Abstract

**Background:** Several studies have been conducted to compare the use DBS as alternative to plasma specimens, but mainly using Whatman 903^®^ cards as filter paper. The aim of this study was to evaluate Whatman FTA^®^ cards (FTA cards) specimens for HIV-1 viral load testing by comparing it to plasma specimens, using 2 real-Time PCR assays.

**Methodology:** A cross-sectional study was conducted between April 2017 and September 2017, in HIV-1 patients admitted at Yalgado Ouédraogo teaching hospital. Paired FTA cards and plasma specimens were collected and analyzed using Abbott RealTi*m*e HIV-1 assay (Abbott) and COBAS^®^ AmpliPrep/COBAS^®^ TaqMan v2.0 (Roche), following manufacturers’ protocol.

**Results:** A total of 107 patients were included. No Statistical differences (p-value > 0.05) were observed between the mean viral loads obtained from FTA cards and plasma specimens with Roche and Abbott assays. Twenty-nine samples with Roche and 15 samples with Abbott assay showed discrepant results. At viral loads of ≤1000 copies/mL, the sensitivity and specificity of FTA cards were 78.6%, and 100% with Roche, and 92.3% and 95.9% with Abbott. Strong correlation was found between FTA cards and plasma specimens with both assays. With Roche, Bland-Altman analysis showed bias of −0.3 and 95% limits of agreement of −2.6 to 1.8 log10, with 97/99 cases (97.9%) within agreement limits. With Abbott, Bland-Altman analysis showed bias of −0.1 and 95% limits of agreement of −2.3 to 2.1 log10, with 96/99 cases (96.9%) within agreement limits.

**Conclusion:** Our study demonstrated the feasibility of using FTA cards filter paper for HIV-1 viral load testing. However, further studies are required for FTA cards filter paper validation in HIV-1 treatment monitoring.

## Introduction

Viral load (VL) testing is the gold standard for HIV treatment monitoring. Periodic VL tests are the most accurate way of determining whether antiretroviral therapy (ART) is working to suppress replication of the virus [1–3]. With ART rapidly expanding in resource-limited settings, VL is a fundamental and even crucial issue in scaling up antiretroviral treatment. However, many barriers exist to VL testing in resource-limited settings, including lack of basic essential equipment, storage and transport limitations for whole blood and plasma[4]. Due to the lability of viral RNA, whole blood in EDTA/K3 tubes cannot be stored more than 6 hours at 25°C[5,6]. Plasma storage and transport require that plasma is transported within 24 hours at 25°C in EDTA/K3 tubes, or within 5 days at 4°C for EDTA/K3 tubes, after centrifugation [7]. In low and middle-income countries, such restrictive guidance on whole blood and plasma transport greatly limits access to VL testing to only those in close proximity to national or regional laboratories. Therefore, a simple method is needed to allow access to HIV-1 VL testing for patients in rural areas.

Since June 2013, WHO has been recommending the use of dried blood spot (DBS) as alternative to plasma for collection, transport, and HIV-1 VL testing and genotyping drug resistance [8,9]. DBS are an inexpensive and practical alternative to plasma; samples are easy to transport, without the need for cold chains or complex equipment; a further benefit of DBS is the reduction in blood sample volume [10,11]. Numerous studies carried out in Burkina Faso and other countries have shown a strong correlation between DBS and plasma specimens for HIV-1 VL testing [12–16]. However, in most of these studies, Whatman 903^®^ cards filter paper was the only filter paper that has been used for HIV-1 load [17]. Diversifying the type of filter papers available for HIV-1 treatment monitoring could reduce shortages risk, decrease costs through price competition and increase the availability of filter paper. In Burkina Faso, another type of paper is now also routinely used for sample collection during malaria vigilance programs and antimalarial drug trials: Whatman FTA^^®^^ cards filter paper.

The aim of this study was to evaluate Whatman FTA^®^ Cards (FTA cards) specimens for HIV-1 VL testing by comparing it to plasma specimens, using COBAS^®^ AmpliPrep/COBAS^®^ TaqMan v2.0 HIV-1 test (Roche) and Abbott RealTime HIV-1 assay (Abbott).

## Methodology

### Study site and design

A cross-sectional study was conducted between April and September 2017, at the National Reference Laboratory for HIV/AIDS and Sexually Transmitted Infections, located at Yalgado Ouédraogo teaching hospital (CHU-YO), 03 BP 7022, Ouaga 03, Ouagadougou, Burkina Faso. Socio-demographic, clinical, and laboratory data were obtained from the study subjects using a structured questionnaire and laboratory analysis blood samples.

### Study population

HIV-1 patients admitted at CHU-YO were the study population. The inclusions criteria were: Patients infected with HIV-1, who consented, antiretroviral-naive patients or patients under antiretroviral treatment.

### Sample collection and processing

Venous collection of whole blood was performed in 2 EDTA/K3 tubes of 4.7 mL or a single EDTA/K3 tube of 10 mL from patients during their routine visits to the CHU-YO. Before plasma separation, DBS were prepared by dispensing 50 μL and 100 μL of blood per spot (2 spots per card) onto FTA cards and dried at room temperature (25 ± 2 C) for 18-24 hours. The FTA cards were stored in zip-lock plastic bags with 2 silica gel desiccants at room temperature upon receipt. Plasma was prepared by centrifugation of the whole blood, aliquoted, and stored at −70 °C until testing for HIV-1 viral load. FTA cards samples were analyzed not more than 14 days after being deposited. Processing of FTA cards for elution of RNA was done as per the protocol for HIV-1 RNA quantitation provided by the ROCHE and ABBOTT diagnostics to be used on their assays systems, COBAS^®^ AmpliPrep/COBAS^®^ TaqMan v2.0 HIV-1 test and m2000rt, respectively.

### Viral Load Quantification

VL was measured from the paired FTA cards and in plasma specimens with Roche and Abbott assays following manufacturers’ protocol. VL results obtained from FTA cards specimens were then compared with that of plasma specimens (gold standard).

### Statistical analysis

The statistical analyzes were performed using RStudio (Version 0.99.903). Sensitivity, specificity predictive positive value and predictive negative value were estimated to determine the performances of FTA cards for quantification of HIV-1 VL at a viral load threshold of 1000 copies/ml, a decision point for therapeutic efficacy. Bland-Altman analysis was used to measure agreement in viral load values obtained from FTA cards and plasma specimens. Correlations between viral loads obtained from FTA cards and plasma specimens were assessed with the Pearson statistical test. All HIV-1 VL values were log10 transformed prior Bland–Altman and correlation analysis. The significance level was set at a p-value of 0.05.

### Ethical considerations

An informed consent form was presented to each patient prior to blood collection and patients who gave verbal consent were enrolled. Additionally, in order to guarantee confidentiality, random anonymous identification numbers were assigned to each patient.

## Results

### Patient characteristics

A total of 107 patients were included. The mean age of the patients was 42.0 ± 13.4 years (ranging from 1 days to 77 years). Most patients were female (sex ratio =0.39).

### Sample collection and bioanalysis

Whole blood was collected from all the patients and the paired FTA cards and plasma specimens collected were analyzed with Abbott and Roche assays for HIV-1 RNA VL. Of all 107 paired FTA cards and plasma specimens tested, 8 FTA cards specimens gave an invalid result with the Abbott assay and were thus excluded from further analysis. Thus, out of 107 paired FTA cards and plasma samples collected, 99 were analysis.

### Comparison between FTA cards and plasma specimens in HIV-1 RNA quantitation

#### *With Roche* assay

No statistical differences (p-value = 0.1704) were observed between the mean VL obtained from FTA cards (1.75 log10) and plasma (1.37 log10) specimens (**Figure 1A**).

**Figure 1:**
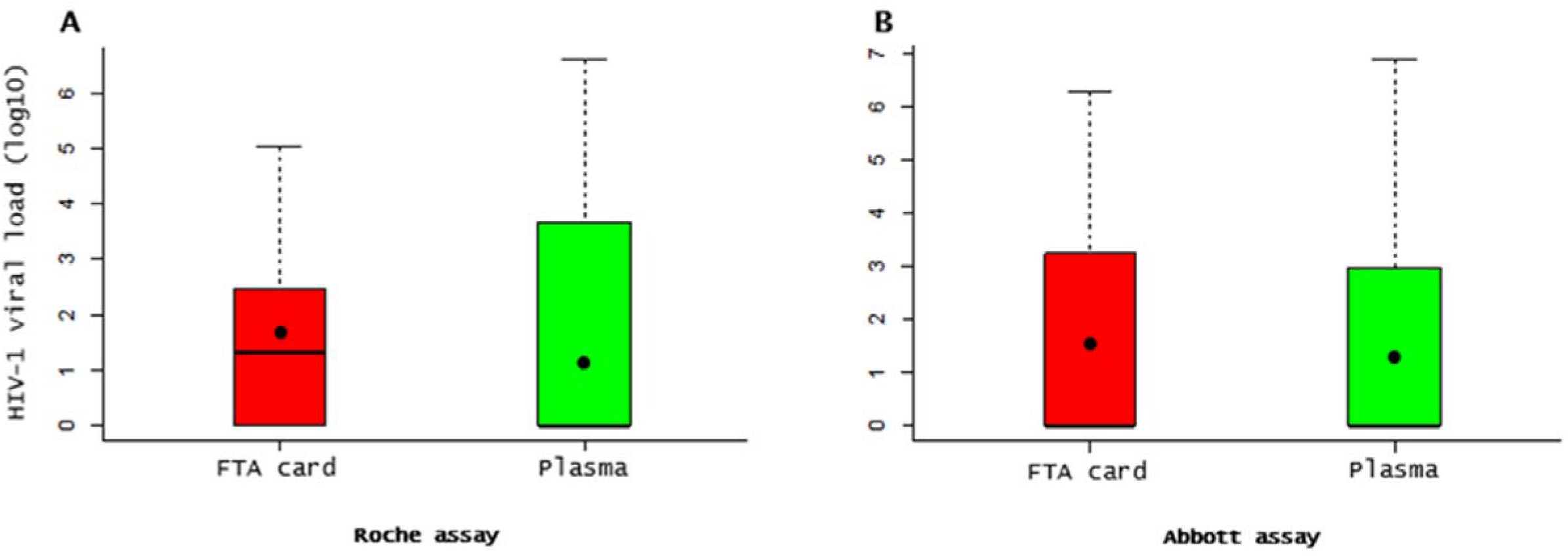
Comparison between Whatman FTA^®^ cards and plasma specimens in HIV-1 RNA. The boxplot to the left **(A)**: using Roche assay; the boxplot to the right **(B)**: using Abbott assay. The black points in the boxplot indicates the means values.

A total of 29 samples showed discrepant results. Eight (17.0%) samples tested not detected on FTA cards but were moderate positive (n=7; 14.9%) and high positive (n=1; 2.1%) on plasma specimens. Twenty-one (70.0%) samples tested moderate positive on FTA cards specimens but gave not detected (n=16, 53.3%) and high positive (n=5; 16.7%) results on plasma specimens (**Table 1**).

**Table 1:**
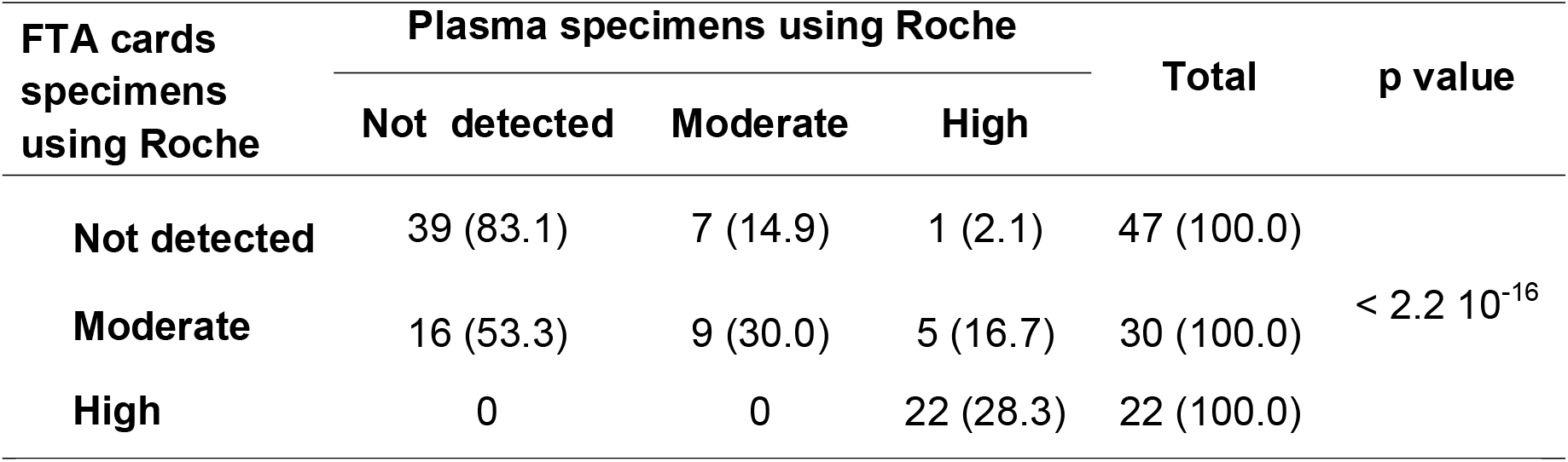
HIV-1 viral load using Whatman FTA^®^ cards and plasma specimens with Roche assay

#### *With Abbott* assay

No statistical differences (p-value = 0.72) were observed between the mean VL obtained from FTA cards (1.50 log10) and plasma (1.38 log10) specimens (**Figure 1B**).

A total of 15 samples showed discrepant results. Twelve (16.7%) samples tested not detected on FTA cards specimens but were moderate positive (n=7; 14.9%) and high positive (n=2; 2.8%) on plasma specimens. Three (11.1%) samples tested high positive on FTA cards specimens but gave not detected (n=1; 3.7%) and moderate positive (n=2; 7.4%) results on plasma specimens (**Table 2**).

**Table 2:**
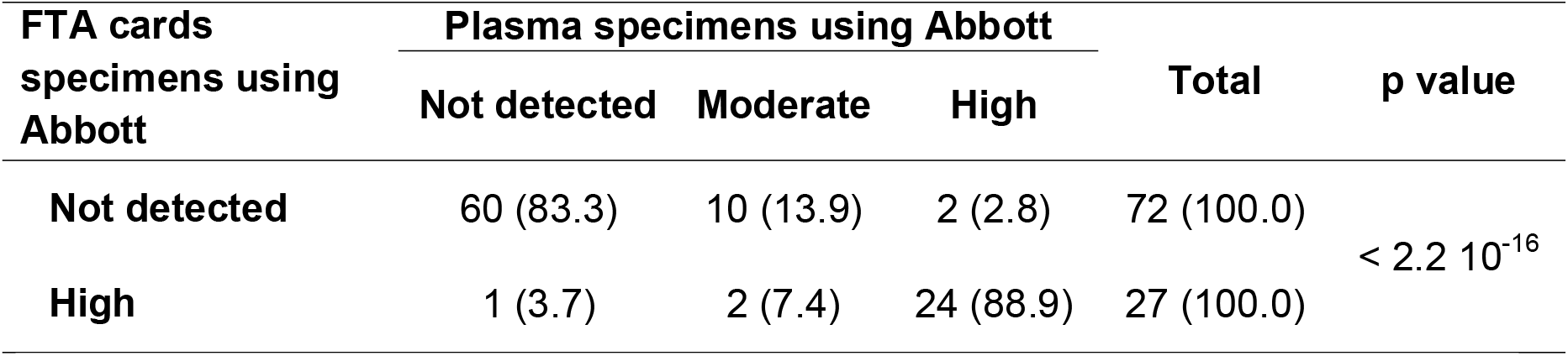
HIV-1 viral load using Whatman FTA^®^ cards and plasma specimens with Abbott assay

| FTA cards specimens using Abbott | Plasma specimens using Abbott |  |  | Total | p value |
| --- | --- | --- | --- | --- | --- |
|  | Not detected | Moderate | High |  |  |
| Not detected | 60 (83.3) | 10 (13.9) | 2 (2.8) | 72 (100.0) | < 2.2 10<sup>-16</sup> |
| High | 1 (3.7) | 2 (7.4) | 24 (88.9) | 27 (100.0) |  |

### Performance of FTA cards for HIV-1 RNA quantitation

#### *With Roche* assay

The sensitivity and specificity of FTA cards at VL of ≤1000 copies/mL were 78.6%, and 100%, respectively (**Table 3**).

**Table 3:**
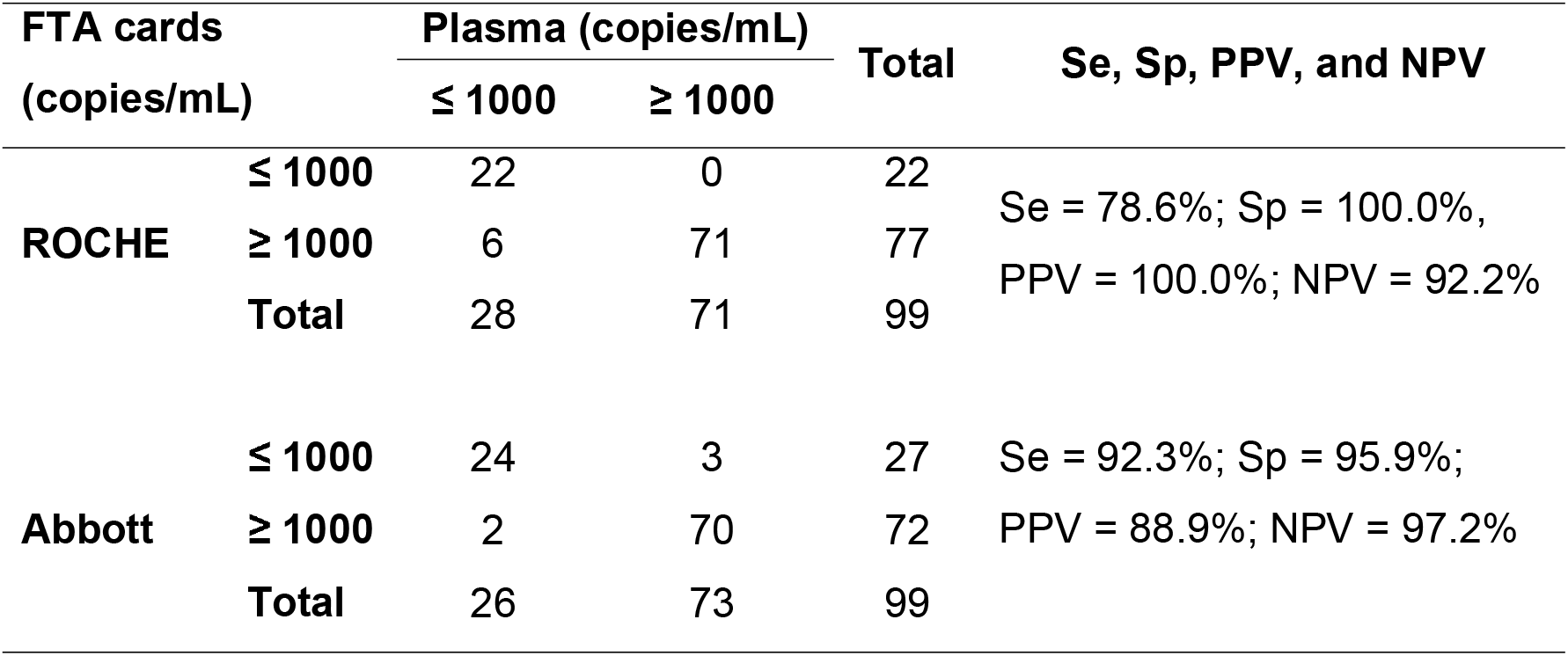
Sensitivity, specificity, positive predictive value, and negative predictive value of Whatman FTA^®^ cards compared with paired plasma specimen for HIV-1 viral load testing at a 1000 copies/mL medical decision pointTable 1: HIV-1 viral load using Whatman FTA^®^ cards and plasma specimens with Roche assay

#### *With Abbott* assay

The sensitivity and specificity of FTA cards at VL of ≤1000 copies/mL were 92.3% and 95.9%, respectively (**Table 3**).

### Correlation and agreement between FTA cards and plasma specimens in HIV-1 RNA quantitation

#### *With Roche* assay

There was a strong correlation (R^2^ = 0.790; p-value < 2.2e-16) between FTA cards and plasma specimens values (Figure 2A). Bland-Altman analysis showed bias of −0. 3 and 95% limits of agreement of −2.6 to 1.8 log10; total number of cases within agreement limits in this study was 97/99 (97.9%) (**Figure 3A**).

**Figure 2:**
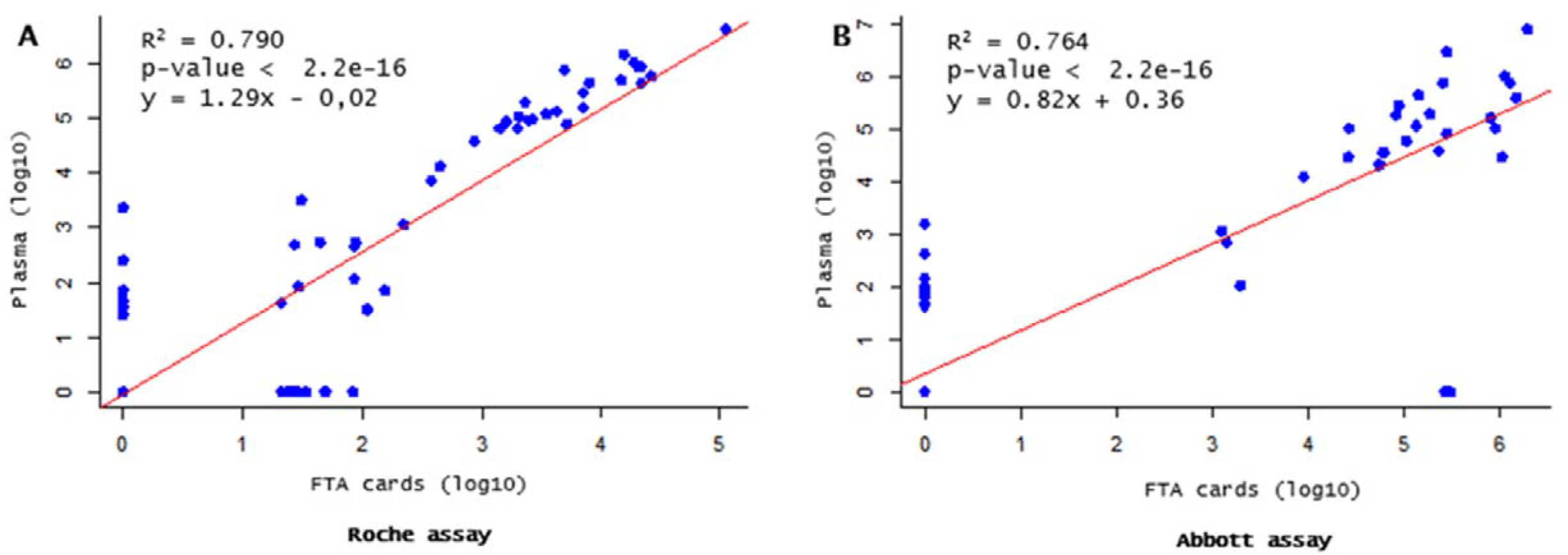
Correlation between FTA cards and plasma specimens in HIV-1 RNA quantitation. A: using Roche assay; B: using Abbott assay.

**Figure 3:**
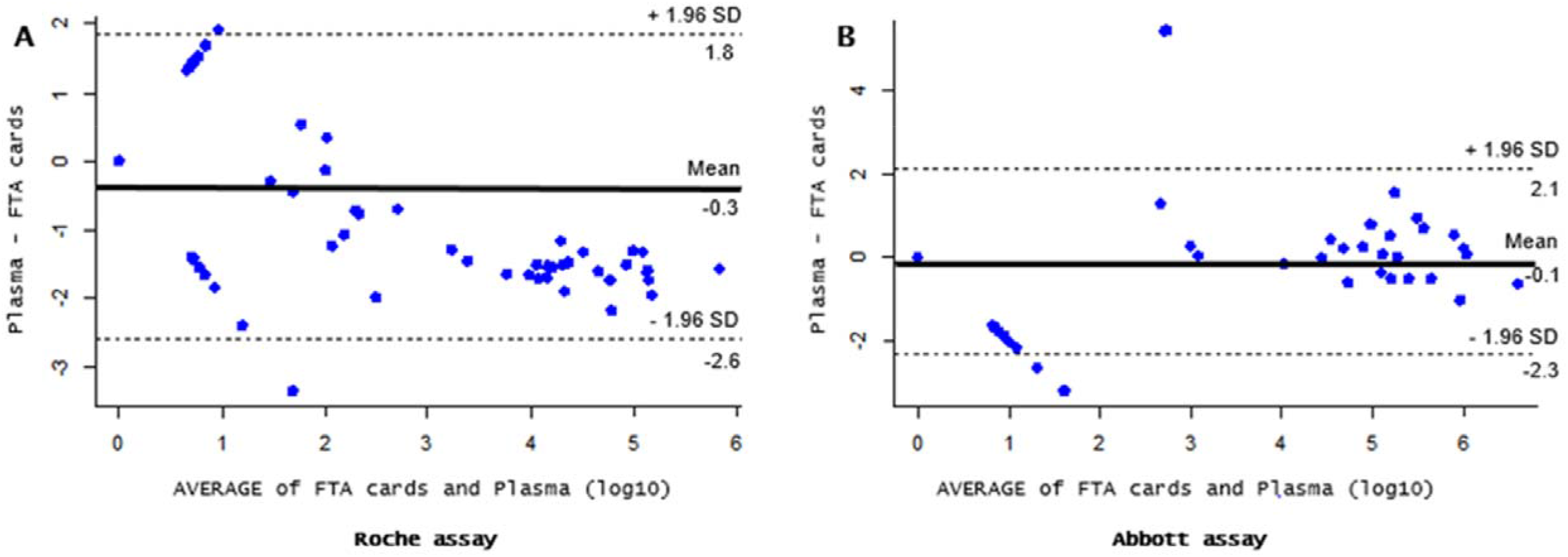
Bland Altman analysis between FTA cards and plasma specimens in HIV-1 RNA quantitation. **A:** using Roche assay; **B:** using Abbott assay.

#### *With Abbott* assay

The correlation between VL values obtained from FTA cards and plasma specimens tested was strong (R^2^ = 0.764; **p-value** < 2.2e-16) (Figure 2B). Bland-Altman analysis showed bias of −0.1 and 95% limits of agreement of −2.3 to 2.1 log10; total number of cases within agreement limits in this study was 96/99 (96.9%) (**Figure 3B**).

## Discussion

Several studies have been conducted to compare the use DBS as alternative to plasma specimens, but mainly using Whatman 903^®^ as filter paper [17]. In Burkina Faso, another type of paper (FTA cards) is now also routinely used for sample collection during malaria vigilance programs and antimalarial drug trials. In this study, FTA cards was evaluated as an alternative sample collection method to plasma for HIV-1 RNA quantitation using commercial Roche and Abbott assays. To the best of our knowledge, this was the first study to evaluate and compare the use of FTA cards filter paper (for DBS) to plasma specimens for VL testing using both Roche and Abbott assays.

In our study, no statistical differences (p-value > 0.05) were observed between the mean VL obtained from FTA cards and plasma specimens using Roche and Abbott assay. These findings are similar to those of previous reports obtained using Whatman 903^®^ cards [18–20]. However, in this study, 17.0% samples tested not detected on FTA cards, were positive on plasma specimens, with 2.1% high positive. This observation of discrepant results is consistent with findings from other studies using Whatman 903^®^ cards [15,21–23]. The reasons of these discrepant results were well documented in the literature[15,17,21–23]. In a systematic review published in 2014, Smit et al. [17] have indicated that the most important reasons DBS is not, and may never be, as sensitive as plasma is because of the differences in sample volume between DBS and plasma. In the current study, the sample volume used on FTA cards was 50 μL and 100 μL, with the Roche and Abbott assays, respectively. Hematocrit has been suggested to recalculate DBS VL to plasma VL copies/mL by applying the difference between plasma and DBS sample volume [24,25]. To make the calculation, hematocrit values can be obtained to adjust DBS VL results by calculating the amount of plasma in a DBS sample. The current study did not used hematocrit adjustment. Indeed, according to the manufacturer’s protocol, a hematocrit adjustment is not required for the calculation of VL obtained from DBS with the Roche and Abbott assays.

An over estimation of HIV-1 RNA levels in FTA cards specimens with low-level viremia (below 1000 copies/mL) was observed in this study; this overestimation was also highlighted in the Bland-Altman analysis (means difference of −0.3 and – 0.1 log10, with Roche and Abbott assays, respectively). This observation is consistent with findings by others with DBS [9,17,19,21,23]. A possible and most advanced explanation of this repeated finding could be the contribution of intracellular HIV-1 DNA and RNA which is present in the DBS but not in the plasma counterpart [26,27]. Vidya et al. [18] suggested that the contribution of intracellular HIV-1 DNA and RNA could be more relevant to specimens with low or undetectable levels viremia than to specimens containing higher levels of extracellular HIV-1 RNA.

At the clinical threshold of 1000 copies/mL, sensibility of FTA cards in this study was seen at 78.6% with Roche, which was slightly less than that observed with Abbott assay (Se = 92.3%) and in already most available literature using Whatman 903^®^ cards [15,21–24,28]. Inversely, our results were higher than that observed in Vietnam using Whatman 903^®^ cards with Roche assay [29]. Additionally, the contribution of HIV cell-associated DNA and RNA could be the reason for slightly lower sensibility for Roche assay. Another possible explanation of the sensitivity observed with Roche could be the elution protocol used in the current study. Following manufacturers’ instructions, time to incubation of DBS is 10 minutes in thermomixer at 1000 rpm, 56°C.

Both Roche and Abbott assays in this study showed a good correlation and agreement between FTA cards and plasma values which is similar to other studies comparing DBS (using Whatman 903^®^ cards) to plasma specimens with Roche and Abbott assays [12,14,15,20].

Our study has some limitations. First, the sample size was restricted. Second, FTA cards were not blotted via finger-prick blood. Third, our study was a laboratory-based study, so the impact of FTA cards sample transport conditions was not evaluated. This study gives preliminary elements to investigations for a longitudinally designed study with a stronger power, incorporating additional factors, such as transport and storage under local conditions, to further evaluate FTA cards specimens for HIV-1 VL testing.

## Conclusion

In summary, we demonstrated the feasibility of using FTA cards for HIV-1 VL testing. FTA cards was found to be a sensitive and specific alternative to plasma testing for HIV-1 VL testing using Abbott assay. Both Roche and Abbott assays showed a good correlation and agreement between FTA cards and plasma values. This information is relevant when considering how to improve access to VL testing by diversifying the type of filter papers available in resource-limited settings.

A study which will increase the testing population size and compare the use of Whatman FTA^®^ to Whatman 903^®^ cards specimens for VL testing using both Roche and Abbott assays is planned for the future.

## Acknowledgments

This research has been supported by the *Programme Sectoriel Santé de la Lutte contre le VIH/Sida et les IST* (PSSLS-IST) through the National Reference Laboratory for HIV/AIDS and Sexually Transmitted Infections. The findings and conclusions in this paper are those of the authors and do not necessarily represent the views of the PSSLS-IST.

## Authors Contributions

L.S. conceived and designed the experiments; A.Y. actively participated to the specimen collection and the study design; A.Y., M.C, G.K.D., H.S., D.C., K.O. performed the experiments. A.Y. analyzed the data; A.Y. and L.S. wrote the manuscript. All authors read and approved the final manuscript.

